# Lipid Bilayer Membranes with Asymmetrically Distributed LPC and DAG

**DOI:** 10.1101/2025.08.13.670206

**Authors:** Chang Liu, Zhongjie Han, Rui Ma, Chen Song

## Abstract

The complex chemical and biophysical characteristics of biomembranes are influenced by the asymmetric distribution of specific lipids. In vitro, the introduction of lysophosphatidylcholine (LPC) into one leaflet of lipid bilayers is frequently utilized to regulate membrane protein activity. In vivo, the conversion of phosphatidylinositol 4,5-bisphosphate (PIP_2_) to diacylglycerol (DAG) in one leaflet can also modulate membrane protein activities. However, the effects of such variations in lipid composition, lipid quantity, and particularly the asymmetry of lipid distribution on the properties and morphology of biomembranes remain to be fully elucidated. Through molecular dynamics simulations, we demonstrate that the asymmetric distribution of LPC and the asymmetric conversion of PIP_2_ induce asymmetric alterations in membrane structures and lipid dynamics. Such alterations can generate an imbalance in the lateral pressure distribution between the two leaflets, potentially leading to membrane curvature. The extent of membrane curvature is also influenced by the length and degree of unsaturation of the lipid acyl tail chain. Our findings underscore the critical role of lipid asymmetry in shaping biomembrane structure and dynamics, providing new insights into the regulation of membrane proteins and cellular functions mediated by these specific lipids.

## 1 Introduction

Biological membranes are semipermeable barriers that compartmentalize cellular and organelle contents. They play a crucial role in facilitating various biological processes, such as regulating the transport of chemical substances, establishing and maintaining transmembrane solute gradients, and modulating intercellular recognition and adhesion. ^1,2^ Biological membranes are primarily composed of lipids, membrane proteins, and carbohydrates. The composition of lipids in biological membranes can vary significantly and is vital to the characteristics of the membrane and its biological functions. ^3,4^ The simplest geometries of biological membranes are planar and spherical. However, membranes can undergo morphological changes across a wide range of length scales, from nanometers to micrometers, and over diverse time scales. ^5–7^ In addition to planar and spherical morphologies, biological membranes can also adopt other long-lived shapes, as observed in the endoplasmic reticulum (ER), the Golgi complex (GC), and the mitochondrial inner membrane (MIM). ^8–10^ In contrast to these persistent membrane architectures that define organelle and cellular structures, transient membrane deformations play essential roles in dynamic processes, including endocytosis, exocytosis, immune responses, cell motility, and division. ^11–13^

Membrane properties are influenced by lipid composition through various physical and chemical characteristics, including head group size, charge, acyl chain length, and degree of saturation. ^14^ Among these characteristics, the relative size of the lipid head group and the hydrophobic tail significantly influences the shape of the membrane. For instance, phosphatidylcholine (PC), the most abundant zwitterionic phospholipid in eukaryotic cell membranes and subcellular organelles, ^15^ possesses a cylindrical shape due to its two acyl chains and large choline head group, which promotes the formation of a planar bilayer. In contrast, lysophosphatidylcholine (LPC), a derivative of PC consisting of a single acyl chain, demonstrates an inverse conical shape that facilitates the formation of inverted hexagonal phases. ^16,17^ On the other hand, diacylglycerol (DAG), which lacks a phosphate head group and thus presents a small polar region, adopts a conical shape that promotes hexagonal phase formation.

These lipid characteristics determine membrane morphology and also regulate the functions of membrane proteins. For instance, LPC, when asymmetrically intercalated into one leaflet of a bilayer membrane, is known to facilitate the gating of mechanosensitive ion channels, such as the bacterial MscL and MscS channels, ^18–22^ as well as the eukaryotic TRAAK and TREK-1 channels. ^23^ Similarly, phosphatidylinositol 4,5-bisphosphate (PIP_2_), a crucial anchoring lipid in the inner leaflet, undergoes hydrolysis by phospholipase C to produce DAG and inositol-1,4,5-triphosphate (IP3), thereby regulating numerous signaling pathways. ^24–27^ This conversion can activate multiple ion channels, including TRPC3, TRPC5, and TRPC6. ^28^ Therefore, alterations in lipid composition can influence not only membrane properties but also the activation of membrane proteins.

Molecular dynamics (MD) simulations are a valuable tool for examining biological membranes, ^29–31^ as they provide a comprehensive atomistic understanding of biological systems when combined with experimental observations. ^30^ However, a detailed atomic-level investigation into the influence of asymmetrically distributed lipid molecules, particularly specific lipids extensively used in experiments such as LPC and DAG, on membrane properties has not been thoroughly explored. Therefore, in this study, we employed all-atom MD simulations to investigate how variations in lipid composition influence membrane characteristics.

We specifically investigate the influence of lipid headgroup and acyl chain structures on membrane properties by symmetrically incorporating 25% LPC of varying lengths into POPC lipid bilayers, or by symmetrically converting 25% PIP_2_ with different saturation levels into DAG (Fig. 1). For the asymmetric systems, the lipid compositional variations were confined to the lower leaflet. Furthermore, we conducted these lipid modifications exclusively in one leaflet of the bilayer to replicate experimental conditions for activating MS channels, allowing us to examine how these asymmetric lipid compositions affect membrane properties. Our results demonstrate that the asymmetrical insertion of LPC thinned the membrane, asymmetrically altered the order of the lipid acyl tail chains in the two leaflets, increased membrane fluidity, and induced curvature toward the LPC-inserted leaflet. Conversely, the asymmetrical conversion of PIP_2_ to DAG thickened the membrane, maintained lipid order with only minor changes, increased fluidity, and caused the membrane to curve away from the DAG leaflet.

**Figure 1:**
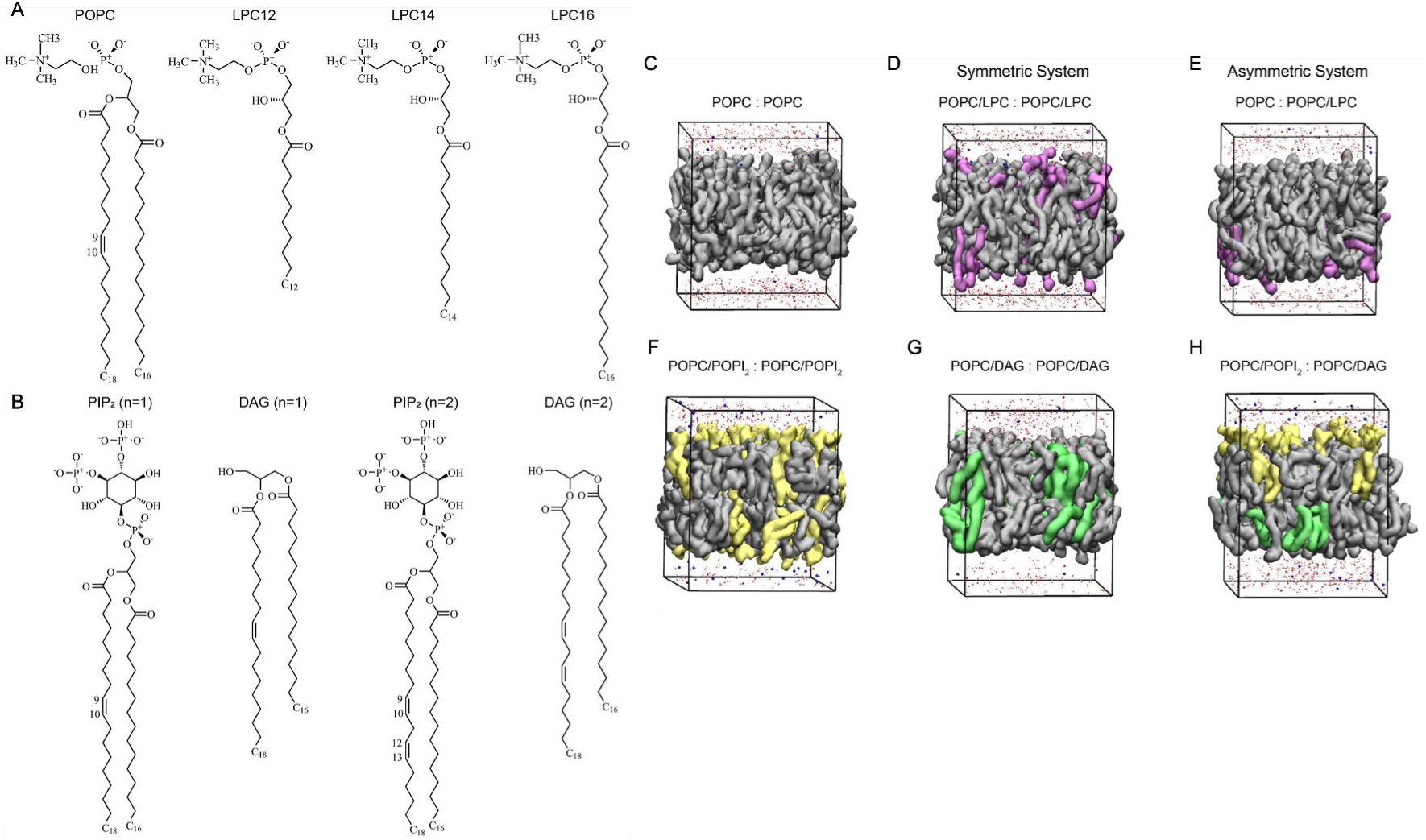
Lipid molecules used in our molecular dynamics (MD) simulations and simulation systems of various lipid compositions. (A) Illustration of POPC and LPC lipids with varying acyl chain lengths. (B) Representation of PIP_2_ or DAG with different degrees of unsaturation in their acyl chains, where *w* denotes the degree of unsaturation. (C) Simulation composed of pure POPC lipids. (D, E) Simulations composed of POPC and asymmetrical or symmetrical LPC. (F, G) Simulations composed of POPC and symmetrical PIP_2_ or DAG. (H) Simulation composed of POPC, asymmetrical PIP_2_, and DAG. POPC lipids are shown with grey surfaces, LPC lipids with pinkish-purple surfaces, PIP_2_ lipids with yellow surfaces, and DAG lipids with green surfaces. For clarity, only part of the ions and water molecules are shown.

## 2 Results

### 2.1 Impact on the overall structure of lipid bilayers

Area per lipid (APL) and membrane thickness are commonly used metrics for assessing the structural properties of lipid bilayers during MD simulations. Membrane expansion, indicated by APL, can be calculated by dividing the total area of the lipid bilayer by the number of lipids present. Membrane thickness is defined as the average distance between the phosphorus atoms in the two leaflets of the lipid bilayer. In this study, we separately calculated the average area occupied by the lipid head (area per lipid head, APLH) and the acyl chain (area per lipid tail, APLT) to gain a clearer understanding of the effects of lipid composition and lipid asymmetry on membrane properties (Fig. 2, Fig. 3). Due to the absence of phosphate groups, DAG was excluded from the calculations of APLH.

**Figure 2:**
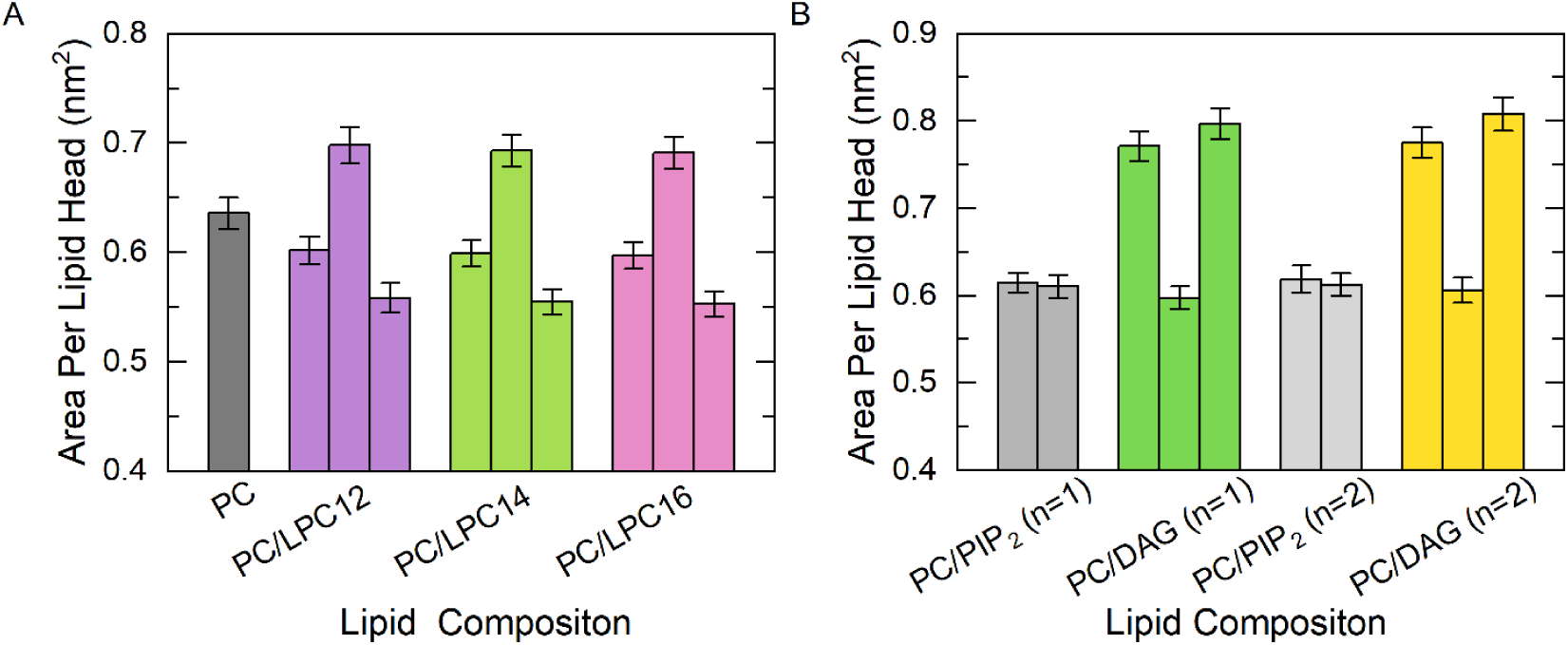
Area per lipid head in LPC insertion (A) and PIP_2_ conversion (B) lipid bilayer systems. In each panel, except for the leftmost histogram, sets of histograms illustrates the results for three different systems: the symmetric system on the left, the upper leaflet system in the middle, and the lower leaflet-altered system on the right. In the symmetric system, when the number of lipid head groups is equal in both the upper and lower leaflets, only one histogram is displayed. The error bars represent the standard deviation obtained from five independent MD trajectories. w denotes the degree of unsaturation.

**Figure 3:**
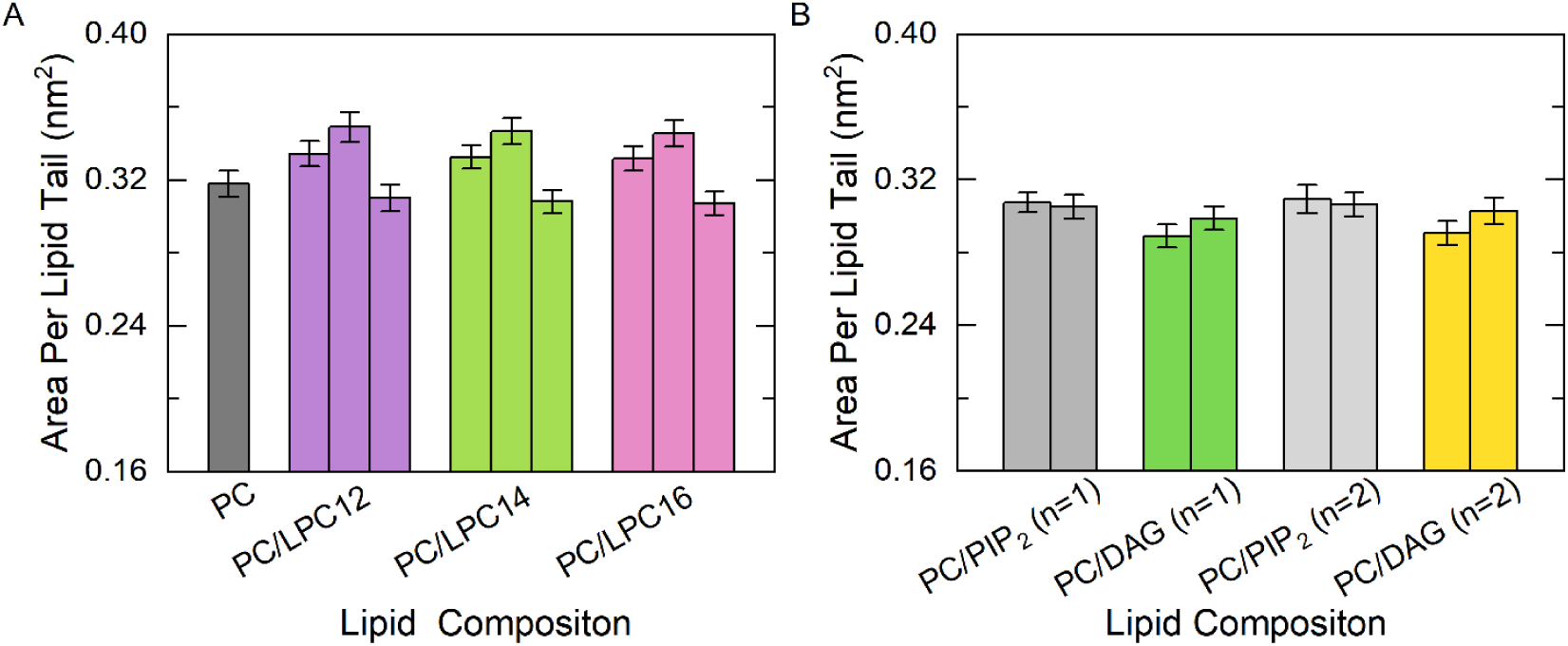
Area per lipid tail in LPC insertion (A) and PIP_2_ conversion (B) lipid bilayer systems. In each panel, except for the leftmost histogram, sets of histograms illustrate the results for three different systems: the symmetric system on the left, the upper leaflet system in the middle, and the lower leaflet-altered system on the right. In the symmetric system, where the number of lipid tails is equal in both the upper and lower leaflets, only one histogram is displayed. The error bars represent the standard deviation calculated from five independent MD trajectories. w denotes the degree of unsaturation.

In symmetrical bilayer systems, the insertion of LPC resulted in an approximately 5.4% decrease in APLH and a 5.1% increase in APLT compared to pure POPC lipid bilayers. Conversely, the conversion of PIP_2_ to DAG led to an approximate 25% increase in APLH and a 5.9% decrease in APLT compared to the POPC/PIP_2_ lipid bilayers (Fig. 2, Fig. 3). In asymmetrical bilayer systems, the insertion of LPC or the conversion of PIP_2_ to DAG in the lower leaflet resulted in varied effects on the upper and lower leaflets of the membrane. When LPC was inserted in the lower leaflet, the APLH of this leaflet decreased by approximately 12.2%. However, it caused an approximate 9.6% increase of APLH in the upper leaflet compared to pure POPC lipid bilayers (Fig. 2A). The APLT increased by around 10% in the upper leaflet and decreased by 2.5% in the lower leaflet (Fig. 3A). When PIP_2_ was converted to DAG in the lower leaflet, the APLH of this leaflet increased by approximately 30%. However, this conversion led to an approximately 2.5% decrease of APLH in the upper leaflet (Fig. 2B). Since DAG lipids share the same tail acyl chain as PIP_2_ lipids, the conversion of PIP_2_ to DAG had a minor effect on the average lipid tail in both symmetrical and asymmetrical bilayer systems (Fig. 3B). The asymmetric bilayer system exhibited a more pronounced impact on the upper leaflet compared to the corresponding symmetric bilayer systems, in the absence of lipid composition changes. The length of the LPC tail chain did not significantly affect the area per phosphorus headgroup and acyl tail chain, while the unsaturation of the PIP_2_ or DAG tail chain had a minor effect (Fig. 2, Fig. 3).

Variations in bilayer thickness were observed following alterations in lipid composition. As the insertion length of the LPC acyl chain tail increased, the reduction in thickness was less pronounced compared to that of pure POPC. Similarly, an increase in the unsaturation of PIP_2_ converted to DAG resulted in a smaller increase in bilayer thickness. LPC insertion resulted in a decrease in bilayer thickness, while PIP_2_ conversion led to an increase. These changes in bilayer thickness were more pronounced in symmetric bilayer systems compared to asymmetric bilayer systems (Fig. 4).

**Figure 4:**
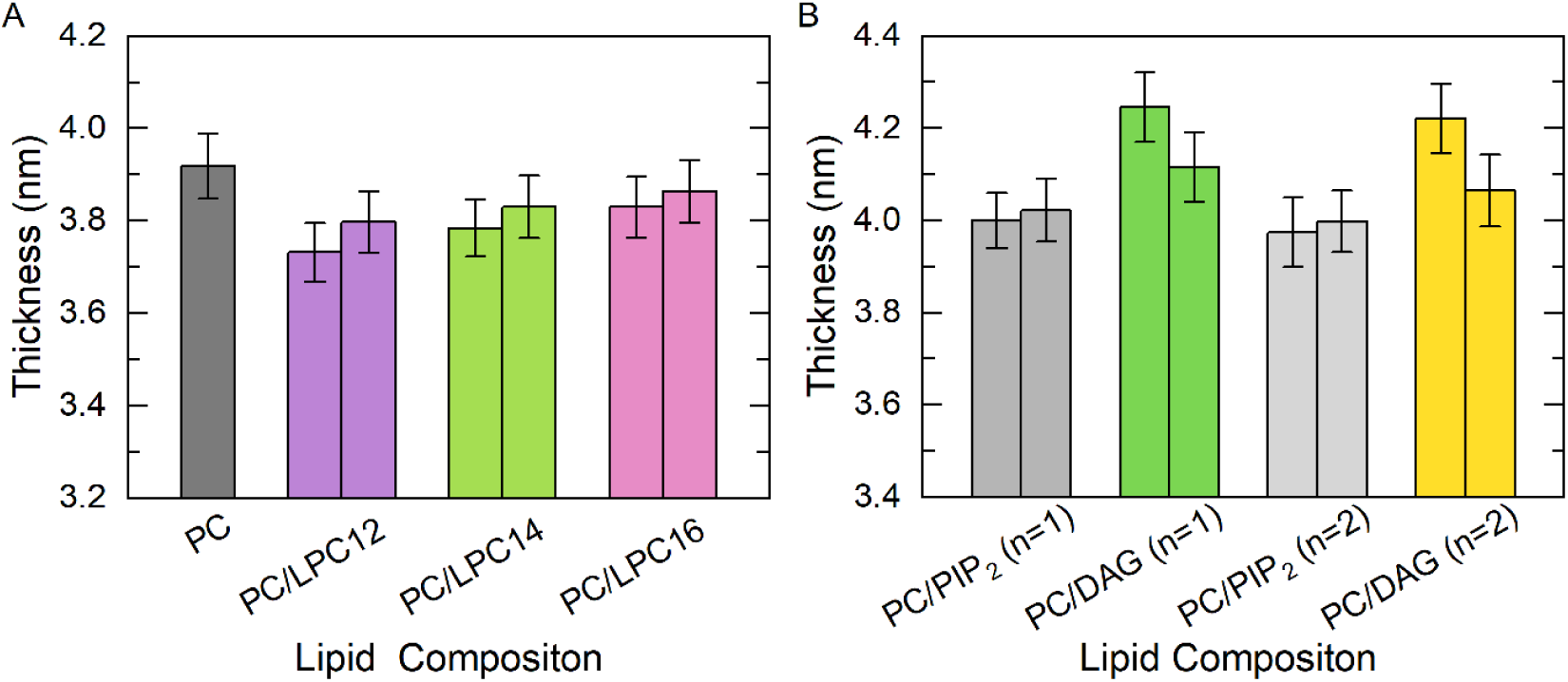
Lipid bilayer thickness in LPC (A) and PIP_2_ conversion (B) systems. The set of histograms illustrates the results for two systems: the symmetric system on the left and the asymmetric system on the right. w denotes the degree of unsaturation.

### 2.2 Impact on the order parameter and fluidity of lipid molecules

The deuterium order parameter (|*S_CD_*|) of lipid tails is a critical indicator of the orientation and ordering of the carbon chains. The order parameters range from 1 to −0.5, where a value of 1 indicates a fully ordered arrangement along the membrane normal, while lower values indicate a progressively disordered distribution that is perpendicular to the membrane normal. ^32^ To evaluate how lipid composition influences the structure of lipid bilayers, we analyzed |S_CD_| of the acyl chains (Eq. (1), Fig. 5, Fig. 6).

**Figure 5:**
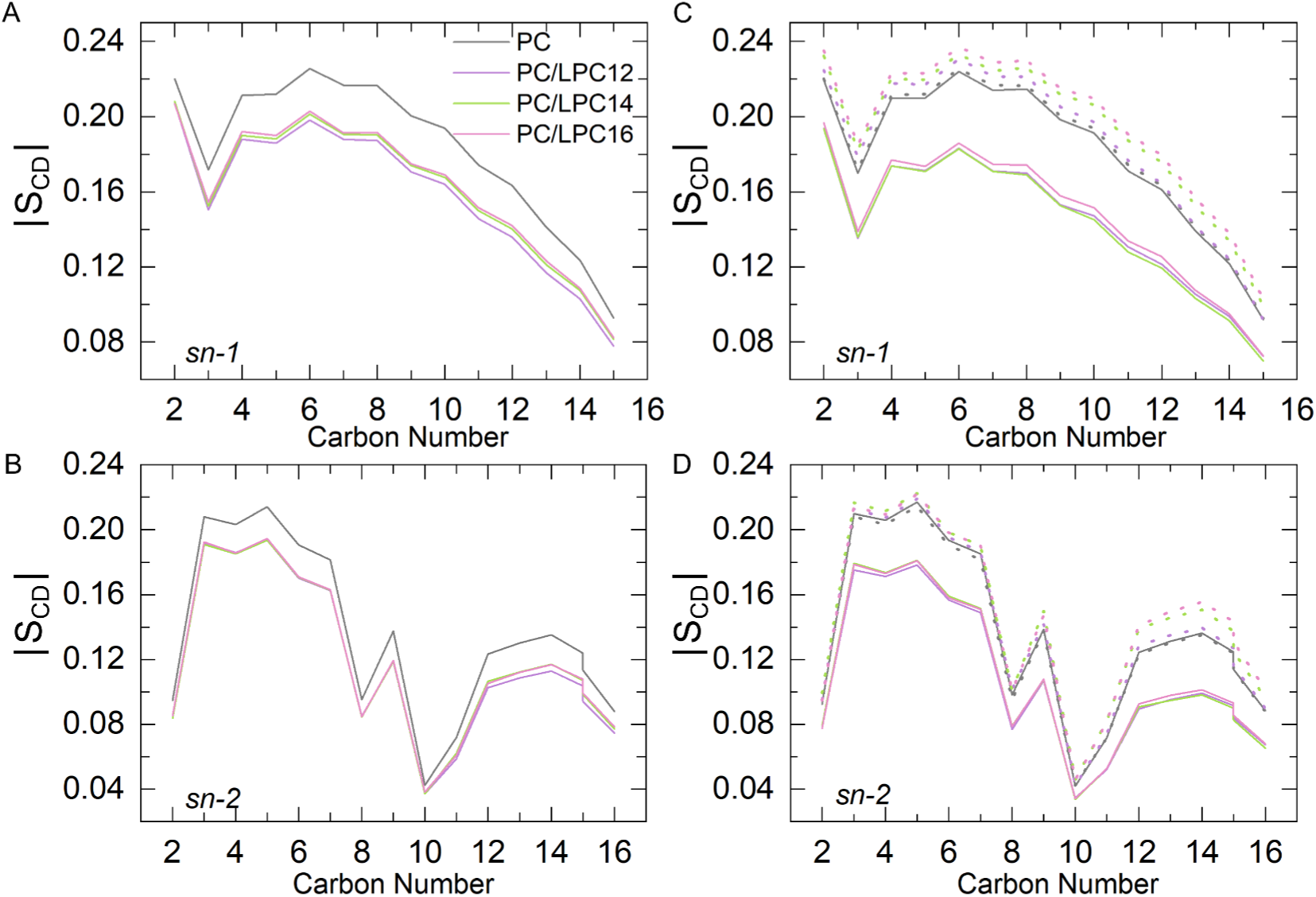
Acyl chain order parameters of POPC lipids in LPC insertion bilayer systems. The order parameters of the two acyl chains of POPC, designated as sn-1 and sn-2, were analyzed in both symmetric (A, B) and asymmetric (C, D) systems. The solid line represents the order parameters for the upper leaflet, while the dotted plot denotes the order parameters for the lower leaflet (C, D). w denotes the degree of unsaturation.

**Figure 6:**
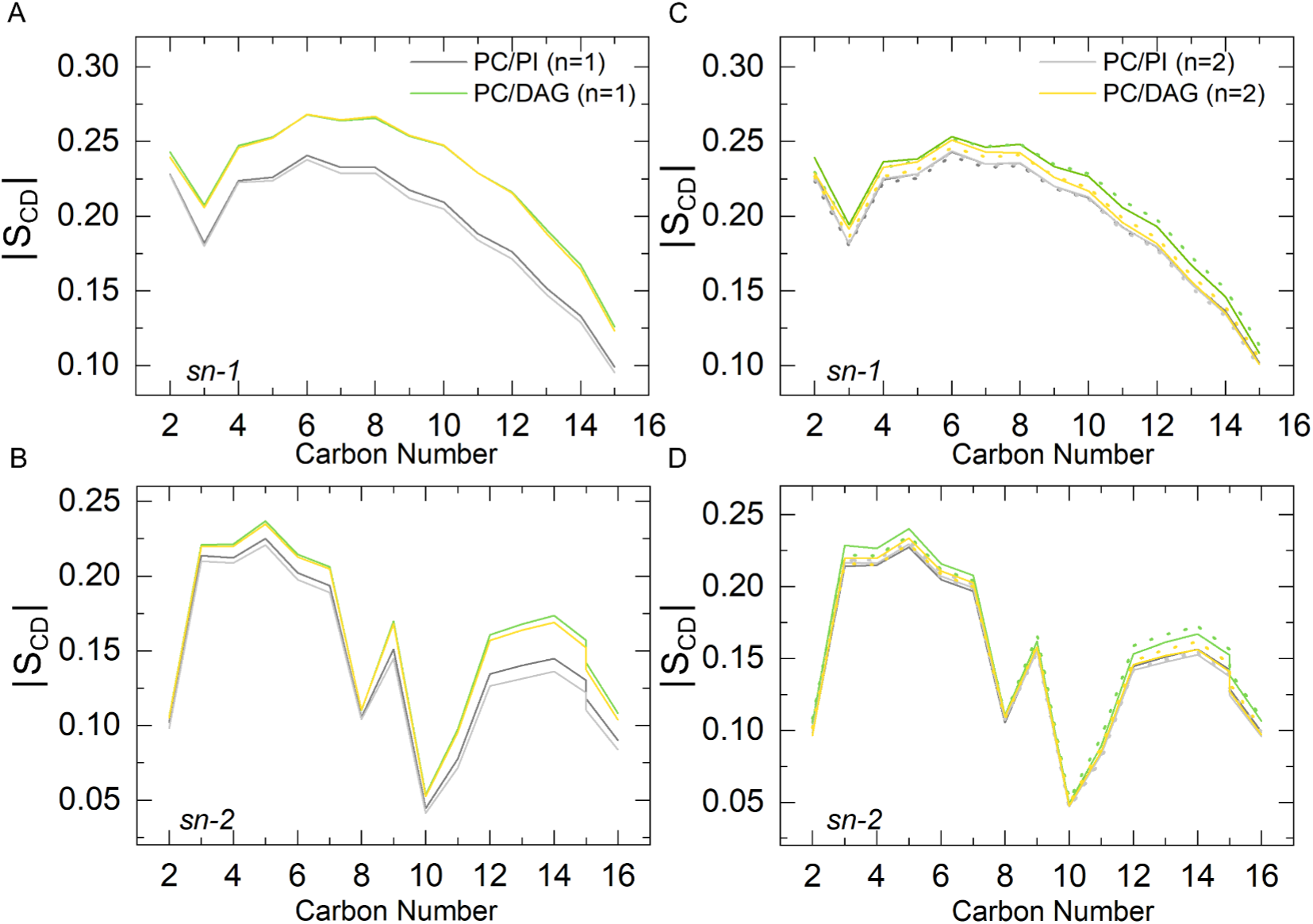
Acyl chain order parameters of POPC lipids in PIP_2_ conversion bilayer systems. The order parameters of the two acyl chains of POPC, designated as sn-1 and sn-2, were analyzed in symmetric (A, B) and asymmetric (C, D) bilayer systems. The solid line represents the order parameters for the upper leaflet, while the dotted plot denotes the order parameters for the lower leaflet (C, D).

In symmetric bilayer systems, LPC insertion resulted in an increase in membrane disorder (Fig. 5A–B), while the conversion of PIP_2_ to DAG led to a slight increase in lipid ordering (Fig. 6A–B). In asymmetric bilayer systems, LPC insertion exhibited distinct effects on the two lipid leaflets. In the lower leaflet where LPCs were inserted, the lipid acyl chains demonstrated a slight increase in ordering, while the upper layer exhibited a greater degree of disorder (Fig. 5C–D). These effects were less pronounced in bilayer systems involving PIP_2_-DAG conversion (Fig. 6C–D). These observed changes were consistent with the results of the APLT analysis. As illustrated in Fig. 3, larger APLT values indicated an increased available space for the lipid acyl chains, which can accommodate a more disordered tail. The order parameters of the POPC acyl chain were not significantly influenced by the length or degree of unsaturation of the lipid acyl chain (Figs. 5 and 6).

The lateral diffusion coefficient of lipids is commonly used to quantify the mobility of lipid molecules within the membrane plane. This parameter is crucial to elucidate the dynamic properties of biological membranes, which play a key role in the regulation of various biological processes, such as lipid phase transitions and clustering of lipids and proteins. ^33,34^ In our atomistic MD simulations, we determined the lateral diffusion coefficients of lipid molecules by analyzing the mean square displacement of lipid phosphorus atoms (Fig. S1). The diffusion coefficient was calculated by fitting a straight line to the data from 100 to 150 ns, as described in Eq. (2), for both symmetric simulation systems and asymmetrical LPC insertion systems. For the PIP_2_-DAG conversion systems, a straight line was fitted to the data from 150 ns to 200 ns. LPC insertion increased the fluidity of the POPC lipid bilayer in both symmetric and asymmetric bilayer systems. The diffusion coefficient of the LPC-inserted leaflet decreased with increasing length of the LPC acyl chain in both symmetric and asymmetric bilayer systems. However, no significant differences were observed in the upper leaflets of the asymmetric bilayer systems (Fig. 7A). The conversion of PIP_2_ to DAG in the symmetric and asymmetric bilayer systems increased the fluidity of the POPC lipid bilayers. In both types of systems, the diffusion coefficient of the leaflet undergoing PIP_2_ conversion increased with the unsaturation of the acyl chain. However, compared to the leaflet where PIP_2_ is converted to DAG, the fluidity of the upper leaflet in the lipid bilayers of asymmetric conversion did not show significant changes (Fig. 7B).

**Figure 7:**
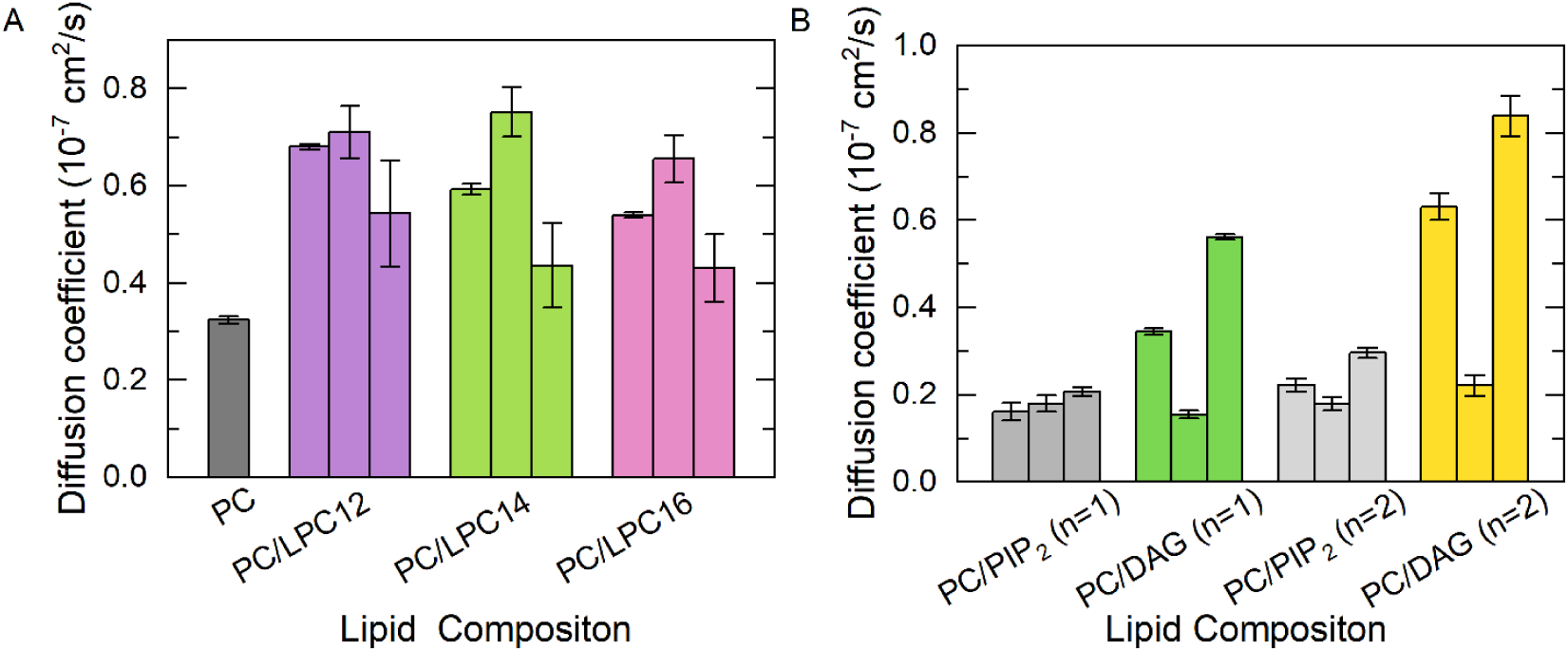
Lateral diffusion coefficients of POPC lipid molecules in symmetric or asymmetric LPC insertion bilayer systems (A) and lateral diffusion coefficients of lipid molecules in symmetric or asymmetric PIP_2_ conversion bilayer systems (B). In each panel, sets of histograms illustrate the results for three distinct systems: the symmetric system on the left, the upper leaflet system in the middle, and the lower leaflet-altered system on the right. In the symmetric system, only one histogram is displayed due to the equal composition in both leaflets. w denotes the degree of unsaturation.

### 2.3 Asymmetrical alteration induced asymmetric pressure profiles across lipid bilayers

To further elucidate the effect of changes in lipid composition on membrane properties, we calculated the lateral pressure profiles (LPPs) along the membrane normal. LPPs depict the spatial distribution of lateral stresses within a lipid bilayer, encompassing repulsive pressure components from lipid-lipid interactions and cohesive hydrophobic tension that promotes segregation of lipid chains from the aqueous environment. These components exhibit an uneven distribution across the bilayer. ^35^ The pressures across the monolayer must be balanced for a lipid monolayer to remain flat. If the lateral pressure of the chain and the head group region is unbalanced, the shape of the monolayer will change. In a symmetric bilayer where the composition and quantity of lipids are identical in both monolayers, they will exhibit the same LPPs and counteract each other. Therefore, we focus on the analysis of LPPs in asymmetric simulation systems (Fig. 8, Fig. 9).

**Figure 8:**
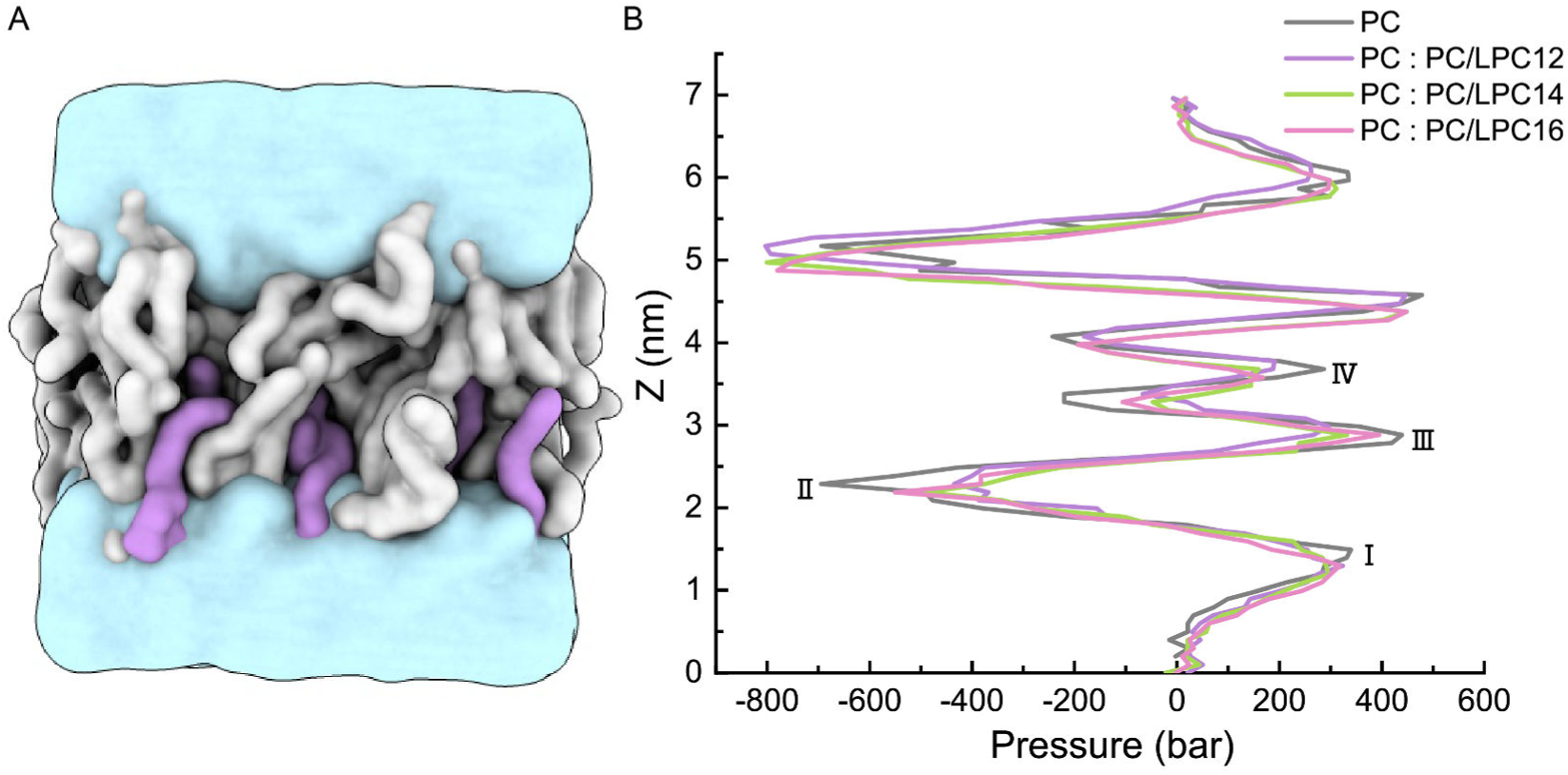
Simulation system of PC : PC/LPC lipid bilayer (A) and lateral pressure profiles in asymmetric LPC insertion systems with varying acyl chain lengths (B).

**Figure 9:**
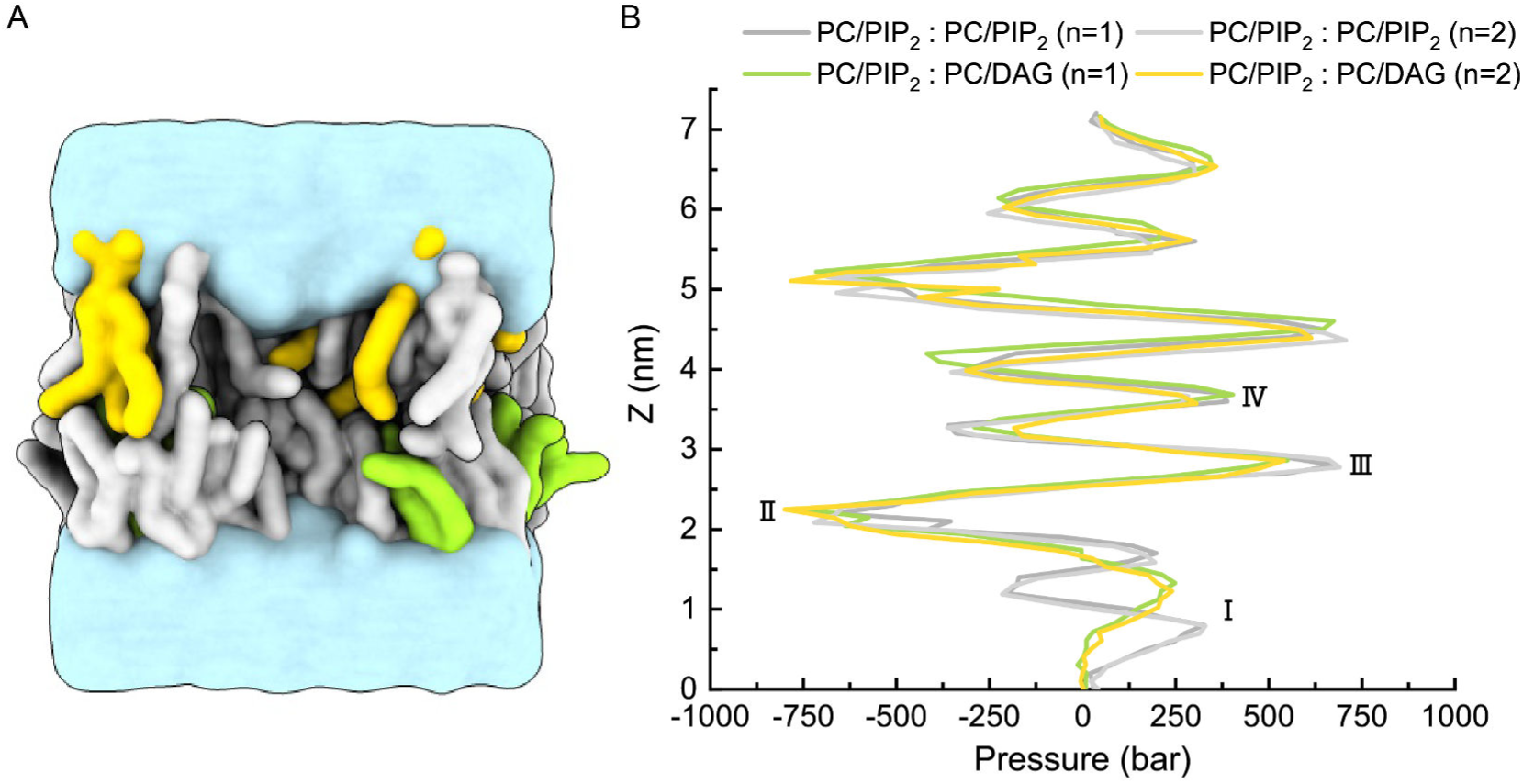
Simulation system of PC/PIP_2_ : PC/DAG lipid bilayer (A) and lateral pressure profiles in asymmetric PIP_2_ conversion to DAG systems with varying degrees of acyl chain unsaturation (B). w denotes the degree of unsaturation.

As can be seen in Fig. 8, Fig. 9, a positive lateral pressure (I) was observed in the headgroup region, primarily due to steric, hydrational, and electrostatic effects. These interactions were overall repulsive, but might incorporate attractive contributions, such as hydrogen bonding interactions. In the vicinity of the glycerol backbone region (II), located just above/below the lipid headgroups, an attractive force arose because of the unfavorable contact of the hydrocarbon chains with water, known as the hydrophobic effect. Within the hydrocarbon interior of the membrane (III), attractive van der Waals interactions between the chains were counteracted by repulsive interactions resulting from the thermal motions of the chains, resulting in a net positive lateral pressure that tends to expand the membrane. Additionally, the lateral pressure profile exhibited a positive peak at the bilayer center (IV), which was related to the geometries of specific lipid types. ^36,37^

In systems with LPC insertion, compared to pure POPC membranes, the attraction in region (II) and the repulsion in region (III) were reduced (Fig. 8). In simulation systems where PIP_2_ converted into DAG (Fig. 9), the repulsion between the headgroups of POPC and PIP_2_ generated two peaks in region (I, grey or light-grey line). And DAG lacks a headgroup, thus exhibiting only one peak resulting from repulsion between the headgroups of POPC lipids (I, green or yellow line).Furthermore, this conversion also diminished the repulsion in the lipid tail double-bond region (III), as evidenced by the weaker peak intensities of the green and yellow curves compared to the gray and light gray ones in Fig. 9.

To analyze the differential stress between the leaflets, we integrated the lateral pressure π(*z*) along the membrane normal *z* direction from the bulk water to the bilayer center (Fig. 8B, Fig. 9B). The differential stress of the two leaflets was then obtained and denoted as Δσ ((σ_lower_) – (σ_upper_)). The mean of Δσ, derived from the analysis of five independently repeated simulation trajectories, was approximately 0 mN/m in pure POPC and POPC : POPC/PIP_2_ (n=1, 2) lipid bilayers (Table 1). This finding suggests that membranes composed of these lipid constituents tend to form planar structures. Interestingly, the lipid composition asymmetry of the POPC and PIP_2_ in our simulation system did not lead to a significant differential stress between the two leaflets.

**Table 1:** Differential stress between two leaflets and corresponding membrane curvature. The error bars represent the standard deviation of Δσ or *c*_0_ obtained from five independent MD trajectories.

| Upper leaflet | POPC | POPC | POPC | POPC | POPC | POPC | POPC | POPC |
| --- | --- | --- | --- | --- | --- | --- | --- | --- |
| Lower leaflet | POPC | POPC/LPC12 | POPC/LPC14 | POPC/LPC16 | POPC/PIP<sub>2</sub> (n=1) | POPC/DAG (n=1) | POPC/PIP<sub>2</sub> (n=2) | POPC/DAG (n=2) |
| $\Delta\sigma$ (mN/m) | $0.46 \pm 0.60$ | $-9.97 \pm 0.76$ | $-11.25 \pm 1.08$ | $-12.83 \pm 0.71$ | $0.84 \pm 0.90$ | $5.56 \pm 1.01$ | $-0.10 \pm 1.2$ | $7.32 \pm 1.74$ |
| $c_0$ (nm<sup>-1</sup>) | $0.01 \pm 0.01$ | $-0.20 \pm 0.01$ | $-0.23 \pm 0.01$ | $-0.23 \pm 0.01$ | $-0.00 \pm 0.01$ | $0.09 \pm 0.02$ | $0.00 \pm 0.01$ | $0.15 \pm 0.02$ |

In contrast, the insertion of LPC and the conversion of PIP_2_ into DAG induced differential stress in the two membrane leaflets in opposite directions. For LPC insertion, the Δσ fluctuated within the range of −9.97 mN/m to −12.83 mN/m, whereas the conversion of PIP_2_ into DAG resulted in the Δσ fluctuating within the range of 5.56 mN/m to 7.32 mN/m (Table 1). This phenomenon can be attributed to LPC having a relatively larger lipid head group and a single-chain acyl chain (Fig. 1A), which leads to increased crowding on the inserted side. Conversely, the conversion of PIP_2_, which has a larger head group, into DAG, which has an almost negligible head group (Fig. 1B), results in a less crowded lipid head group region in the lower leaflet. Moreover, as the length of the LPC acyl chain inserted into the lipid increased and the unsaturation of the acyl chain converted from PIP_2_ to DAG increased, the differential stress became more pronounced (Table 1). These results indicate that both the acyl chain length and the degree of unsaturation of the lipid tails significantly affect the LPPs of the lipid bilayers.

### 2.4 Asymmetric pressure profiles induced membrane curvature

The distribution of lateral pressures and tensions across lipid monolayers and bilayers is essential for determining their spontaneous curvature. The insertion of LPC and the conversion of PIP_2_ to DAG introduce asymmetrical stress within the bilayer, potentially leading to curvature. This tendency for the membrane to bend can be characterized by the spontaneous curvature *c*_0_.

We estimated the spontaneous curvature induced by lipid changes via Eq. (5) based on the integral of the first moment of LPPs obtained from MD simulation analyses. The direction of membrane curvature was defined from the perspective of an observer located below the lower leaflet of the bilayer, looking towards the upper leaflet. A concave curvature of the membrane was defined as positive, while a convex curvature was defined as negative. As shown in Table 1, the spontaneous curvature of POPC and POPC : POPC/PIP_2_ was approximately 0 nm^−1^. The insertion of LPC and the conversion of PIP_2_ to DAG resulted in membrane curvature in opposite directions. Specifically, LPC insertion into the lower leaflet induced negative membrane curvature, ranging from −0.20 nm^−1^ to −0.23 nm^−1^), while the conversion of PIP_2_ to DAG in the lower leaflet induced positive membrane curvature, ranging from 0.09 nm^−1^ to 0.15 nm^−1^. Furthermore, as the LPC tail chain lengthened and the unsaturation of DAG increased, the absolute value of the spontaneous curvature induced by these changes became more pronounced.

## 3 Discussion

The composition of the lipid bilayer, in which membrane proteins are embedded, has been demonstrated to play a crucial role in regulating protein function. The lipid bilayer’s composition influences membrane protein function not only through specific biochemical interactions and lipid binding, such as the electrostatic attraction between polybasic motifs in proteins and negatively charged lipid headgroups (e.g., PIP_2_), ^38^ but also through the physical properties of membranes themselves. ^39–41^ Our results indicate that changes in lipid composition, caused by LPC insertion or PIP_2_-DAG conversion, can significantly impact membrane properties, including APL, thickness, fluidity, lateral pressure, and potentially curvature. These alterations in membrane properties can affect membrane morphology and regulate the activities of membrane-embedded or associated proteins, such as mechanosensitive ion channels.

The LPC insertion and PIP_2_-DAG conversion exhibit distinct effects on APLH, indicating that the packing within the lipid headgroup region is influenced by both the structure of the lipid fatty acyl chain and the organization of the lipid headgroup itself. The hydrophobic thickness of the lipid bilayer is anticipated to closely match that of any protein embedded within it. This alignment is crucial due to the significant energetic penalty associated with exposing either fatty acyl chains or hydrophobic amino acids to an aqueous environment. Any disparity in hydrophobic thickness between the lipid bilayer and the protein is likely to result in distortion of either the lipid bilayer, the protein, or both, in order to minimize this mismatch. In cases of extreme hydrophobic mismatch, a membrane protein may be excluded from the lipid bilayer, as observed in simple model transmembrane α-helices. ^42^ Moreover, extreme mismatch could also induce the formation of non-bilayer phases by the lipids, particularly at low molar ratios of lipid to protein. ^42–44^

The insertion of LPC or the conversion of PIP_2_ to DAG influences membrane thickness. Understanding the impact of changes in lipid composition on membrane thickness could yield valuable insights for experimental studies. For instance, alterations in membrane thickness can regulate the gating of mechanosensitive channels such as MscL, ^18^ MscS,^45^ and TREK-2, ^46^ all of which are known to be sensitive to membrane thickness. Additionally, MscS and PIEZO channels are affected by the degrees of lipid unsaturation, ^45,47,48^ likely due to changes in membrane thickness or fluidity. Remarkably, similar lipid remodeling occurs physiologically during T-cell activation, where DAG generation coupled with PUFA enrichment creates a combinatorial lipid signature that potentially coordinates PIEZO-mediated calcium signaling in immune responses.^49^ This highlights the critical role of membrane lipid composition in modulating immune cell function and mechanotransduction pathways.

The transition from the liquid crystalline phase to the gel and fluid phases induces a pronounced shift in the physical properties of a lipid bilayer, which can significantly influence the activities of membrane proteins, such as the clustering of these proteins. The symmetric or asymmetric insertion of LPC into the membrane lipid layer has distinct effects on the order parameters of the two leaflets of the lipid bilayers. Nonetheless, both insertion modes of LPCs contribute to an increase in membrane fluidity. The conversion of PIP_2_ to DAG did not significantly affect the membrane order parameter but had a substantial impact on membrane fluidity. Alterations in these lipid components manifest effects on membrane order parameters and dynamics, offering a method for studying the conformation of membrane proteins, for example, Ca^2+^, Na^+^, and K^+^-ATPases. ^44,50,51^

Lipid lateral pressure profiles play a crucial role in the conformational equilibrium of proteins, constituting a vital aspect of protein function. ^52^ Mechanosensitive channels, known for their responsiveness to membrane forces and curvature, ^53^ can be activated by membrane stretch, lateral pressure, and local membrane curvature, as well as by the incorporation of amphipathic lipids. ^54^ This activation could be closely linked to the intricate relationship between mechanosensitive channels and membrane lipids. Amphipathic compounds, such as chlorpromazine, local anesthetics, and LPC could induce channel activation through insertion^18,19,55^ or the conversion of PIP_2_ into DAG in one monolayer of the lipid bilayers. ^28^ Whether this activation was triggered by local membrane curvature or changes in the transbilayer pressure profile without local curvature remained unknown. ^56–58^

Membrane tension and bending not only drive gating transitions in mechanosensitive channels but also impact the organization of membrane proteins. ^59^ Certain proteins exhibit a propensity to partition into membrane regions with specific curvature, while others demonstrate flexibility, adapting to a range of curvature states. ^60,61^ The lipid structures of LPC and DAG are not inclined to form planar bilayer structures; instead, they induce membrane curvature. This property can be leveraged in *in vitro* experiments to regulate the localization of specific proteins. For instance, the spatial distribution of PIEZO1 channels, influenced by membrane curvature, avoids localization at highly curved membrane protrusions such as filopodia and instead is enriched in membrane invaginations. ^62–64^

In summary, this study examines the effects of symmetric and asymmetric LPC insertion, as well as the conversion of PIP_2_ to DAG, on the structure, dynamic properties, and morphology of membranes. The findings contribute to a more comprehensive understanding of the roles of LPC and DAG in regulating membrane structures and the functions of membrane proteins. This work not only enriches our insight into related physiological processes but also serves as a valuable reference for future investigations.

## 4 Method

### 4.1 Simulation systems

The molecular dynamics simulation systems consisted of symmetric or asymmetric mixtures of POPC/LPC, POPC/PIP_2_, or POPC/DAG lipid monolayers, as well as pure POPC lipids. The LPC lipids insertion system consisted of 144 lipids and approximately 6417 water molecules. The PIP_2_ conversion system was composed of 128 lipids and approximately 5297 water molecules (Table 2). Na^+^ and Cl^−^ ions were added to the system to achieve charge neutrality and a physiological ion concentration of 0.15 M.

**Table 2:**
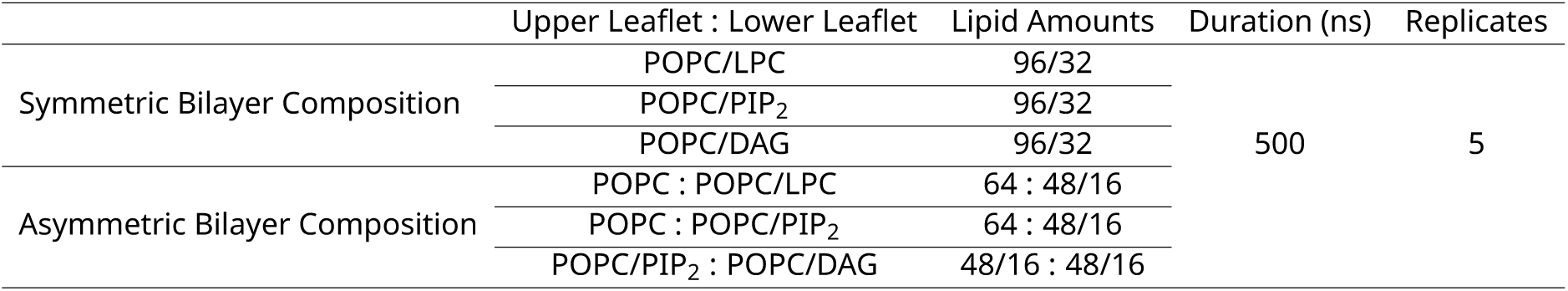
List of the simulation systems.

|  | Upper Leaflet : Lower Leaflet | Lipid Amounts | Duration (ns) | Replicates |
| --- | --- | --- | --- | --- |
| Symmetric Bilayer Composition | POPC/LPC | 96/32 | 500 | 5 |
|  | POPC/PIP<sub>2</sub> | 96/32 |  |  |
|  | POPC/DAG | 96/32 |  |  |
| Asymmetric Bilayer Composition | POPC : POPC/LPC | 64 : 48/16 |  |  |
|  | POPC : POPC/PIP<sub>2</sub> | 64 : 48/16 |  |  |
|  | POPC/PIP<sub>2</sub> : POPC/DAG | 48/16 : 48/16 |  |  |

### 4.2 Molecular dynamics simulations

All MD simulations were performed using GROMACS 2018.6. ^65^ The simulations utilized the CHARMM36 force field and the CHARMM implementation of the TIP3 water model. ^66–68^ The temperature was maintained at a constant temperature of 300 K using the Nose-Hoover thermostat. ^69,70^ The pressure was maintained at 1 bar using the Parinello-Rahman semiisotropic barostat^71,72^ with a time constant of 1 ps and a compressibility of 4.5 × 10^−5 −1^. A cutoff of 1.2 nm was used for the Lennard-Jones interaction, with a smooth shift function from 1.0 to 1.2 nm. The particle mesh Ewald (PME) method was used for long-range electrostatics. ^73^ After energy minimization, we ran 200-ps NVT and 500-ps NPT simulations to preliminarily equilibrate the systems, and then we carried out 500-ns production simulations for each lipid simulation system (Table 2).

### 4.3 Analysis of MD trajectories

If not otherwise stated, the last 400 ns trajectories were used for the analysis. The order parameters of the lipid tails were analyzed using the following definition:

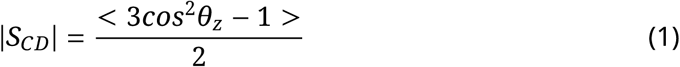

where θ*_z_* is the angle between the C–H bond and the membrane normal (z direction).

The lateral diffusion coefficient of the lipids was calculated with the following equation:

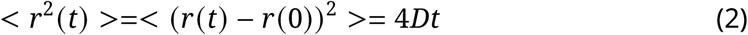

where < *r*^2^(*t*) > is the two-dimensional mean squared displacement (MSD) of phosphorus atoms, D is the diffusion coefficient, and *t* is the simulation time. A time interval of 150-200 ns or 100-150 ns was used to perform a linear fit and obtain the value of D in PIP_2_ conversion systems and other systems. The error estimate was performed by calculating the difference in the diffusion coefficients obtained from the two halves of the fit interval, as implemented in the GROMACS-msd tool.

The lateral pressure profiles (LPPs) were calculated using the GROMACS-LS tool. ^74^ The calculation of LPPs was based on the last 400 ns of the simulation data (Table 2), with the coordinates and velocities saved every 20 ps. An electrostatic cutoff of 2.0 nm was used for the LPP calculation. ^74^ The profiles were calculated with about 80 slabs, corresponding to an approximate slab width of 1 Å. The outputs of GROMACS-LS were the components of the stress tensor, σ, as a function of the *z* coordinate. LPP was calculated using the formula

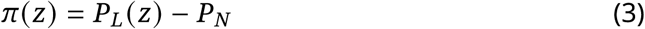

where *P_L_*(*z*) is the lateral component of the pressure tensor (*P_L_*(*z*) = 0.5[*P_XX_* (*z*) + *P_YY_* (*z*)]), and *P_N_* is the normal component (*P_N_* = *P_ZZ_*). The spontaneous curvature estimation was calculated using continuum mechanics theory. When two lipid monolayers were combined to form a bilayer, we used the midplane between the two monolayers to represent the shape of the bilayer, whose-position was denoted as *z* = 0 along the normal direction of the bilayer. Assuming that the thickness of the two monolayers was *d*^+^ and *d*^−^, we calculated the spontaneous curvature of each monolayer via the integral ^75,76^

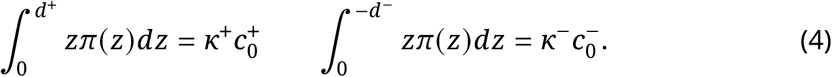

where *k*^±^ and *c*_0_^±^ denoted the bending rigidity and spontaneous curvature of the two monalayers, respectively, and π(*z*) was the LPP defined in (3). In practice, we performed integration from the midplane of the bilayer to bulk water. The spontaneous curvature of the bilayer then reads as

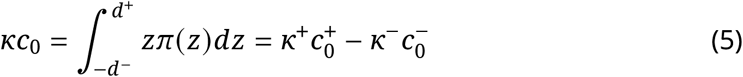

For both the bilayer and the monolayer, the first moment of the LLP π(*z*) gave the product of the bending rigidity and the spontaneous curvature. In order to obtain the spontaneous curvature, the values for the bending rigidity *κ* and *κ*^±^ were needed which in principle could be obtained by a buckling simulation. ^77^ Here, for simplicity, we assumed *κ*^±^ = 10*k_B_T* and *κ* = *κ*^+^ + *κ*^−^ = 20*k_B_T*, which were typical values for the plasma membrane. ^78^ We neglected the effect that the bilayer bending rigidity *κ* could be different for the two monolayers due to compositional asymmetry. This may affect the quantitative accuracy of the calculated spontaneous curvature, but not the sign of it (bending direction of the membranes).

## 6 Acknowledgements

This work was supported by the Innovative Research Groups of the National Natural Science Foundation of China (T2321001 to C.S.) and the Postdoctoral Fellowship of the Peking-Tsinghua Center for Life Sciences (Z.H.). C.S. was supported in part by the Frontier Innovation Fund of Peking University Chengdu Academy for Advanced Interdisciplinary Biotechnologies. Part of the computation was performed on the computing platform of the Center for Life Sciences at Peking University.

## Supplementary Figure

**Figure S1:**
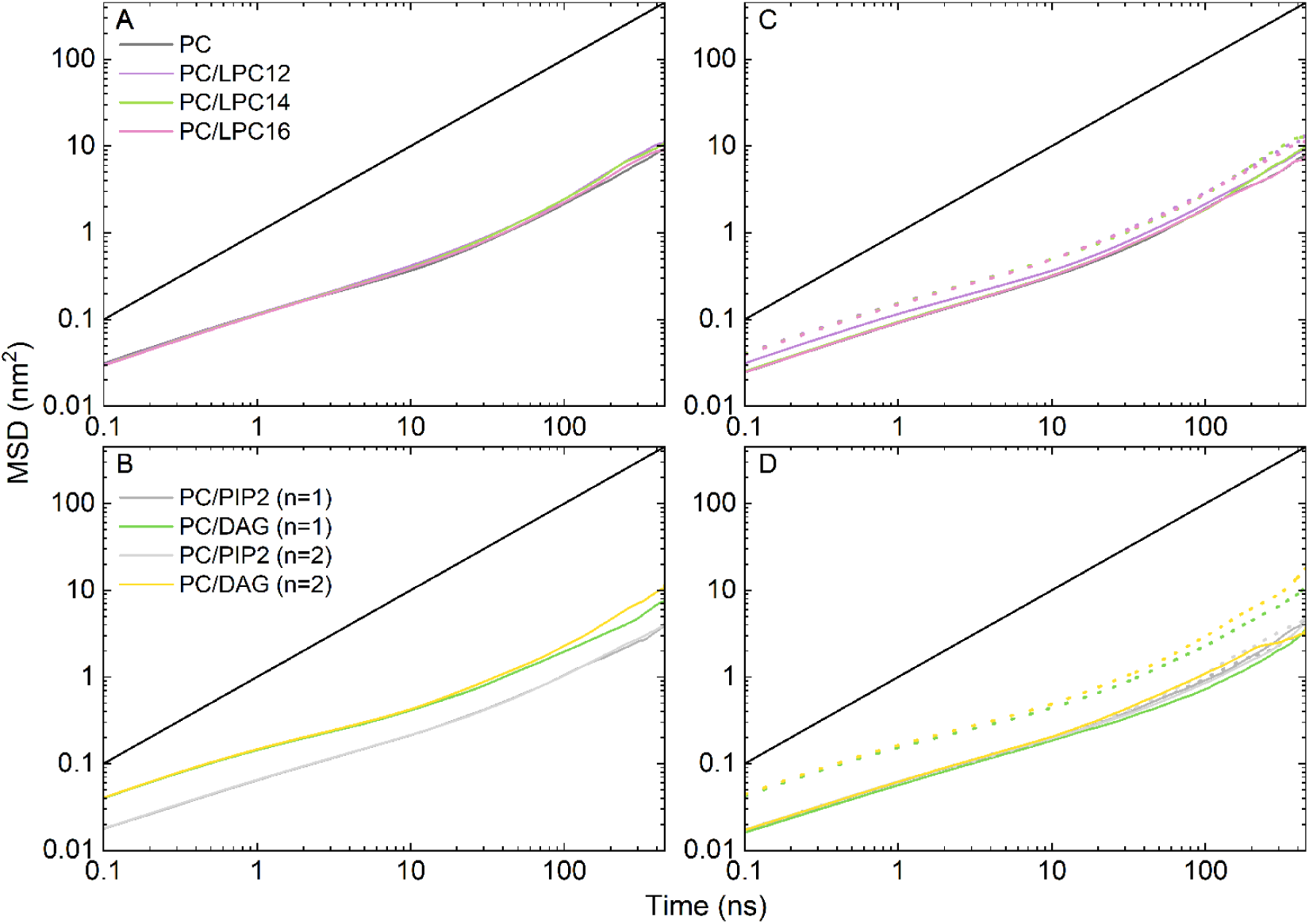
Two-dimensional mean square displacement (MSD) of lipid molecules in bilayer systems with symmetric LPC insertion and PIP_2_-DAG conversion (A, B), or with asymmetric LPC insertion and PIP_2_-DAG conversion (C, D). The solid and dotted lines in panels (C) and (D) represent the MSD of lipid molecules in the upper and lower leaflets, respectively.

